# Arabidopsis O-fucosyltransferase SPINDLY regulates root hair patterning independently of gibberellin signalling

**DOI:** 10.1101/2020.04.23.057083

**Authors:** Krishna Vasant Mutanwad, Isabella Zangl, Doris Lucyshyn

## Abstract

Root hairs are able to sense soil composition and play an important role for water and nutrient uptake. In *Arabidopsis thaliana*, root hairs are distributed in the epidermis in a specific pattern, regularly alternating with non-root hair cells in continuous cell files. This patterning is regulated by internal factors such as a number of hormones, as well as external factors like nutrient availability. Thus, root-hair patterning is an excellent model for studying the plasticity of cell fate determination in response to environmental changes. Here, we report that loss-of-function mutants in the Protein O-Fucosyltransferase SPINDLY (SPY) form ectopic root hairs. Using a number of transcriptional reporters, we show that patterning in *spy-22* is affected upstream of the central regulators GLABRA2 (GL2) and WEREWOLF (WER). O-fucosylation of nuclear and cytosolic proteins is an important post-translational modification that is still not very well understood. So far, SPY is best characterized for its role in gibberellin signalling via fucosylation of the growth-repressing DELLA protein REPRESSOR OF GA (RGA). Our data suggest that the formation of ectopic root hairs in *spy-22* is independent of RGA and gibberellin signalling.

## Introduction

Post translational modifications (PTM) dynamically modulate various physiological and morphological events throughout the life span of plants (Millar *et al.* 2019). O-Glycosylation of nuclear and cytosolic proteins is one such PTM, and plants carry two O-glycosyltransferases responsible for these modifications: the Protein O-Fucosyltransferase (POFUT) SPINDLY (SPY), and the O-GlcNAc Transferase (OGT) SECRET AGENT (SEC) (Hartweck *et al.* 2002; Olszewski *et al.* 2010; Zentella *et al.* 2016; Zentella *et al.* 2017). These proteins regulate significant events in plants, from embryo development to the determination of flowering time and flower development (Hartweck *et al.* 2002; Hartweck *et al.* 2006). *spy* mutants were initially identified due to their resistance to the gibberellin (GA) biosynthesis inhibitor paclobutrazol, leading to constitutively active GA signalling (Jacobsen and Olszewski, 1993; Swain and Olszewski, 1996). Further studies reported that SPY and SEC are involved in GA signalling via modification of the growth repressing DELLA protein RGA (REPRESSOR OF GA) (Silverstone *et al.* 2007; Zentella *et al.* 2016; Zentella *et al.* 2017). *spy* mutants display various phenotypic traits, such as early flowering, early phase transitions, partial male sterility, abnormal trichome formation and disordered phyllotaxy (Silverstone *et al.* 2007). Recently, SEC also was reported to be involved in delaying flowering time in Arabidopsis (Xing *et al.* 2018). The majority of the studies thus have focused on the role of O-glycosylation in aerial tissue development and the subsequent phenotypes are often attributed to its participation in GA signalling. SEC and SPY are also active in roots, however their impact on root development and morphogenesis is largely unexplored (Hartweck *et al.* 2006; Silverstone *et al.* 2007; Swain *et al.* 2002).

Tissue morphology and cellular organisation are decisive for root development in *Arabidopsis thaliana*. Epidermal tissue is comprised of hair-forming trichoblast cells and non-hair-forming atrichoblast cells (Dolan *et al.* 1993; Löfke *et al.* 2015; Scheres and Wolkenfelt, 1998). The arrangement of the hair and non-hair cells is established around the single ring-like layer of cortex cells. A hair cell arises at the junction between and is connected to two cortical cells, while a non-hair cell is usually adhered to only a single cortex cell. Moreover, hair cells are generally separated by non-hair cells between them (Balcerowicz *et al.* 2015; Dolan *et al.* 1994; Salazar-Henao *et al.* 2016). Various transcription factors like GLABRA2 (GL2), WEREWOLF (WER) and CAPRICE (CPC) are responsible for determination of epidermal cell patterning in Arabidopsis. GL2 and WER regulate the establishment of non-hair cells (Lee and Schiefelbein, 1999; Masucci *et al.* 1996), whereas CPC activity is required for the formation of hair cells (Wada *et al.* 1997). GL2 expression is promoted by WER via the formation of a multiprotein complex comprised of TRANSPARENT TESTA GLABRA (TTG1), GLABRA3 (GL3) and ENHANCER OF GLABRA3 (EGL3) (Bernhardt *et al.* 2003; Schiefelbein *et al.* 2014). Further, GL2 establishes non-hair cell fate by supressing the expression of root hair-promoting basic Helix-Loop-Helix (bHLH) transcription factors like ROOT HAIR DEFECTIVE 6 (RHD6), RHD6-LIKE1 (RSL1), RSL2, Lj-RHL1-LIKE1 (LRL1), and LRL2 (Balcerowicz *et al.* 2015; Masucci and Schiefelbein, 1996). On the contrary, in root hair cells, expression of WER is strongly reduced. This allows CPC or its paralogs ENHANCER OF TRY AND CPC 1 (ETC1), ETC3 or TRYPTICHON (TRY) to take its place in the TTG1/EGL3/GL3 complex, resulting in negative regulation of GL2 and de-repression of root hair promoting genes, thus establishing root hair cell fate (Lee and Schiefelbein, 2002; Salazar-Henao *et al.* 2016).

Root hair development is dynamically controlled by environmental factors like reactive oxygen species (ROS) and pH (Monshausen *et al.* 2007). Furthermore, availability of mineral nutrients like inorganic phosphate (Pi) and iron (Fe) in the surroundings also modulates the development and morphology of root hairs (Janes *et al.* 2018; Müller and Schmidt. 2004; Salazar-Henao *et al.* 2016). Similarly, phytohormones like auxin, ethylene and brassinosteroids are known to influence root hair patterning and development (Balcerowicz *et al.* 2015; Borassi *et al.* 2020; Kuppusamy *et al.* 2009; Liu *et al.* 2018; Shibata and Sugimoto, 2019). However, a role of gibberellin (GA) in epidermis morphology, root hair formation and development has not been described as yet, nor a potential role of the O-glycosyltransferases SPY and SEC in this context. *spy* mutants have been previously reported to display an extra layer of cortex cells, the middle cortex (MC), a phenotype associated with high level ROS signalling (Cui *et al.* 2014; Cui and Benfey, 2009). Beyond this, root tissue morphology of *spy* and *sec* mutants is largely unexplored. Hence, we initiated the investigation of the role of SPY and SEC in root development and tissue patterning, also in relation to GA signalling. Here, we show that epidermis morphology and root hair patterning is altered in *spy*, but not in *sec* mutants. Using a set of reporter constructs, we established that SPY regulates patterning upstream of WER. However, we did not find any evidence for an involvement of GA signalling, indicating that SPY regulates root hair patterning independently of DELLA proteins and GA-signalling.

## Results

### The Arabidopsis Protein O-fucosyltransferase mutant *spy-22* has larger root apical meristems

In order to investigate the involvement of O-glycosylation in Arabidopsis root development we analysed various morphological phenotypes of the T-DNA insertion lines *spy-22* and *sec-5* in comparison to wild type Col-0. SPY and SEC regulate GA signalling by modifying the DELLA protein RGA (Silverstone *et al.* 2007; Zentella *et al.* 2016; Zentella *et al.* 2017) and *spy*-mutants display constitutive GA-signalling phenotypes (Jacobsen and Olszewski, 1993). GA deficient mutants like *ga1-3* are reported to have a reduced root apical meristem (RAM) size (Achard *et al.* 2009). To analyse if O-glycosylation is involved in GA-dependent regulation of RAM size, we measured the RAM of 7-day old seedlings, as the region from quiescent centre till the uppermost first cortical cell which is twice as long as wide (Feraru *et al.* 2019). We observed that *spy-22* mutants displayed a significantly longer meristem (347.6 +/- 34.65 µm) compared to the wildtype Col-0 (283.6 +/- 31.92 µm) and *sec-5* (282.4 +/- 27.51 µm) (Figure 1 A, B). On counting the number of epidermal cells in the meristem, we found that the number of cells correlated with meristem size, showing a higher number of cells in *spy-22* (39.10 +/- 4.599) compared to Col-0 (29.05 +/- 3.965) and *sec-5* (28.92 +/- 5.008) (Figure S 1). This result is in line with the effect of increased GA-signalling on cell division and meristem size (Achard *et al*., 2009).

**Figure 1.**
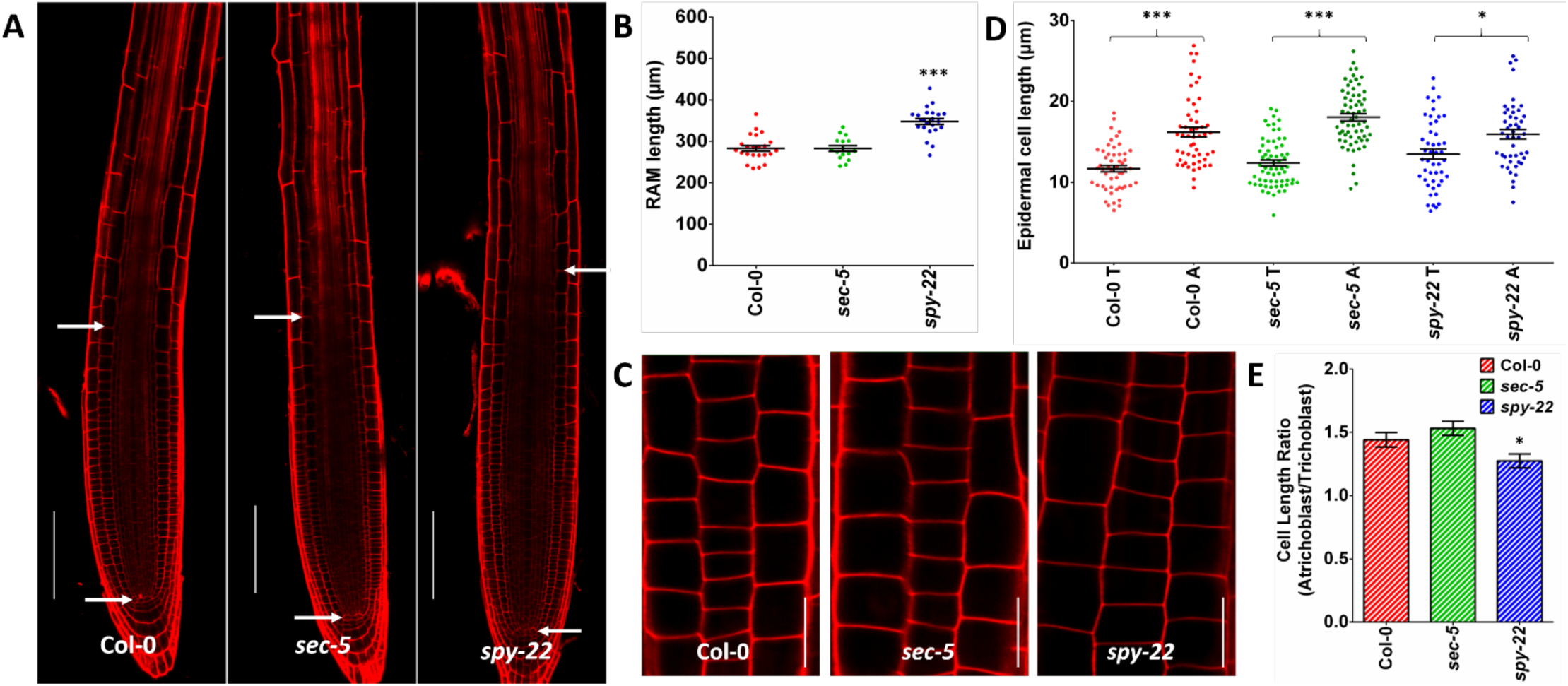
A- Longitudinal cross section images of 7-day old seedlings mounted in PI. Meristem size was defined as the distance from the quiescent center to first uppermost cortical cell which was twice as long as wide, as indicated by white arrows, scale bar – 100µm. B- *spy-22* roots display a significantly longer meristem compared to Col-0 and *sec-5*. n = 16-23. C- The epidermal layer in the late meristematic region of 7-day old seedlings mounted in PI. Lengths of 4 consecutive cells in neighboring (tricho/atrichoblast) files in the late meristem were measured, scale bar – 20µm. D- Atricho- and trichoblast cell length in Col-0, *sec-5* and *spy-22*, n = 47-64. E- The ratio of the epidermal cell lengths of atrichoblasts/trichoblasts is lower in *spy-22* compared to *sec-5* and Col-0. For statistical analysis, One-way ANOVA with Tukey’s multiple comparison and students t-test were done (*** P ≤ 0.001, * P ≤ 0.05), data from three independent biological repeats is shown.

Additional to cell number, also the patterning and distribution of atrichoblasts (non-hair) and trichoblast (hair) cells of the epidermis is crucial in determining the size of the meristematic region in Arabidopsis (Löfke *et al.* 2013). While analysing our mutants, we observed that the difference between atricho- and trichoblast cell sizes was reduced in *spy-22* mutants compared to wild-type and *sec-5*. To quantify that, we measured the lengths of the last four consecutive cells in adjacent (trichoblast and atrichoblast) cell files in the epidermis marking the transition to the root meristem differentiation zone (Lofke *et al.* 2015). We noted that the atrichoblast cells in Col-0 and *sec-5* (16.21 +/- 4.30 µm and 18.05 +/- 3.62 µm respectively) were significantly longer than trichoblast cells (11.70 +/- 2.81 µm and 12.38 +/- 2.95 µm respectively). In *spy-22*, atrichoblast cells (15.92 +/- 4.08 µm) were only slightly longer than cells in corresponding trichoblast files (13.49 +/- 4.30 µm) (Figure 1 C, D). This difference was clearly reflected in a lower ratio of atrichoblast/trichoblast cell length in *spy-22* (1.27) compared to Col-0 (1.44) and *sec-5* (1.53) (Figure 1 E). Taken together, we observed both an increase in cell number, as well as an altered distribution of atrichoblast/trichoblast cell length in *spy-22*, resulting in an increase of root meristem size.

### *spy* mutants display ectopic root hairs

The atypical atricho-to trichoblast morphology in *spy-22* led us to explore the consequences of this observation on root hair development in fully differentiated epidermis cells. In *spy-22*, we frequently observed appearance of two trichoblast cell files developing root hairs adjacent to each other, indicating ectopic root hair formation, while in Col-0 and *sec-5* root hair cell files were always separated from each other by a non-hair cell file (Figure 2 A). The underlying cause for the appearance of ectopic root hairs in *spy-22* was further analysed with the help of reporter lines. We used cell type specific promoter-YFP fusions as described (Marquès-Bueno *et al.* 2016) to monitor the expression of transcription factors implicated in root hair patterning at different stages of development. We initially targeted WER which is involved at an early stage of non-hair cell determination and is expressed strongly in atrichoblast cells and weakly in trichoblasts (Lee and Schiefelbein, 1999). On crossing the WER::4xYFP reporter with *spy-22* and *sec-5*, we observed an uneven signal distribution within single cell files in *spy-22* (Figure 2 B). We also crossed our lines to GL2::4xYFP, which in the wild type is exclusively expressed in the atrichoblasts in the cell division and transition zone. While in Col-0 and *sec-5* a regular pattern of reporter gene expression was observed, GL2 expression in *spy-22* was very patchy, potentially underlying the formation of ectopic trichoblasts within non-hair cell files and vice versa (Figure 2 C). We next employed a reporter that is active in differentiated root hair cells, to determine if expression patterns in the meristematic and transition zone match the patterning of developed root hairs in the differentiation zone. EXP7 is expressed specifically in root hair cells. In EXP7::4xYFP *spy-22* we observed non-hair cells without signal within YFP-positive root hair cell files and vice versa, an aberration in reporter expression which we did not detect in the Col-0 or *sec-5* background (Figure 2 D). Taken together, crosses with various transcriptional reporter lines suggest that SPY regulates root hair patterning upstream of WER.

**Figure 2.**
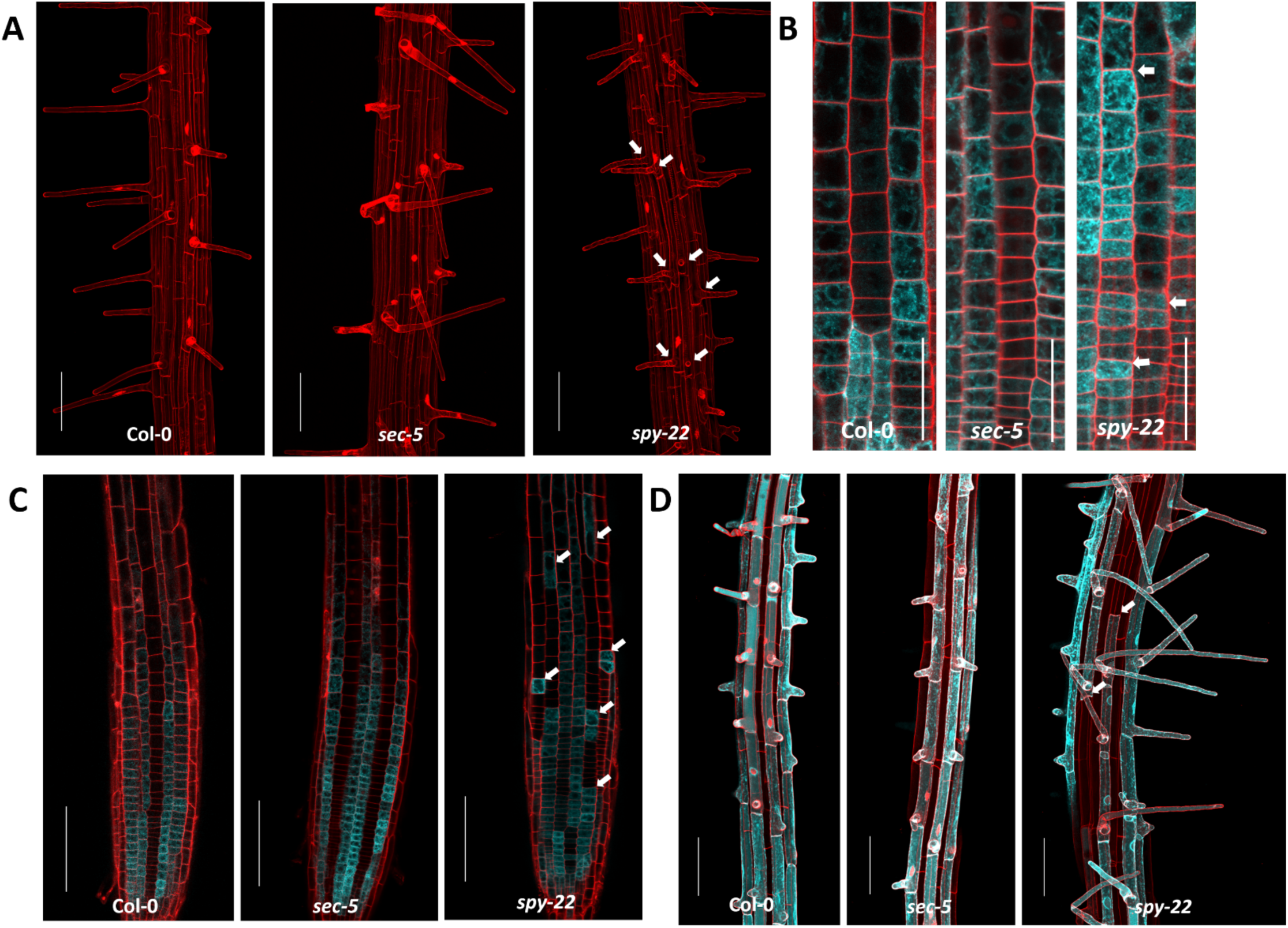
A- Maximum projection of Z stacks to visualize root hair patterning of O-glycosylation mutants. *spy-22* displays ectopic root hairs, scale bar – 100µm. B- WER::4xYFP expression in the epidermal cells in the meristem region. YFP signal in *spy-22* is unevenly distributed within the same cell file, scale bar – 50 µm C- GL2 activity visualized in atrichoblasts expressing GL2::4xYFP. Expression in *spy-22* indicates the presence of trichoblast cells in the atrichoblast cell file and vice versa, scale bar – 100µm. D- EXP7 is exclusively expressed in root hair cells. YFP signal indicates EXP7 promoter activity is not uniform within cell files in *spy-22*, suggesting the presence of atrichoblasts in a trichoblast cell file and vice versa, scale bar – 100µm. Representative pictures of three biological repeats are shown.

It was previously shown that *spy*-mutants generate an additional layer of root cortex cells, which has been attributed to constitutively increased ROS signalling (Cui *et al.* 2014; Cui and Benfey, 2009). This middle cortex between the cortex and the endodermis was also clearly visible in *spy-22* (Figure S2 A). When crossing our lines with SCR::4xYFP to visualize specifically the endodermis, we could confirm the increase in middle cortex formation and clearly distinguish ectopic cell file formation from the endodermis, like seen before (Cui and Benfey, 2009), but there is no indication for a defect in endodermis formation in *spy-22* (Figure S2 B).

### Epidermal cell patterning and ectopic root hair formation in *spy-22* is independent of gibberellin signalling

So far, the best-characterised target of SPY is the DELLA protein RGA, which undergoes a conformational change upon O-fucosylation that enhances the interaction with downstream transcription factors, thereby inhibiting their binding to DNA (Zentella *et al.* 2017). As a result, *spy* mutants show constitutively active GA signalling. So far, GA signalling has not been described to play a role in epidermal cell patterning in *Arabidopsis thaliana*, hence we aimed to understand whether the epidermal patterning of *spy-22* was influenced by increased GA signalling. For initial experiments we treated *spy-22, sec-5* and Col-0 with 10µM GA_3_ and measured the tricho– and atrichoblast cell length in the root meristem transition zone. The distribution pattern remained similar to untreated seedlings, as reported in Figure 1 C. The difference in length of trichoblast cells (13.60 +/- 4.21 µm) and atrichoblast cells (16.15 +/- 3.38 µm) was smaller in *spy-22* when compared to Col-0 and *sec-5* (Figure 3 A), with a lower atrichoblast/trichoblast ratio (1.3) in *spy-22* also after GA_3_ treatment (Figure 3 B), at a ratio comparable to the untreated seedlings (compare Figures 1 E and 3 B). Next, we determined GL2::4xYFP expression in Col-0, *spy-22* and *sec-5* upon treatment with 10 µM GA_3_ and analysed the cell file patterning in the cell division and transition zones. We quantified this phenotype by counting the number of patterning defects (which we defined as the appearance of atrichoblast cells in trichoblast cell files and vice versa) per seedling (Figure 3 C). We observed that Col-0 displayed on average 1.47 patterning defects per seedling, with 7/19 seedlings showing no patterning defects. After treatment with 10 µM GA_3_, frequencies of patterning defects did not significantly change, with an average of 2 per seedling (Figure 3 D). Similarly, there was no significant change in patterning defects in GL2::4xYFP *sec-5* in untreated controls (2.7 patterning defects per seedling) compared to 10µM GA_3_-treated seedlings (2.6 patterning defects per seedling) (Figure 3 D). GL2::4xYFP *spy-22* displayed the highest number of patterning defects per seedling (8.1 per seedling) and this did not change significantly upon treatment with 10 µM GA_3_ (7.6 patterning defects per seedling). These results suggest that exogenous application of gibberellin does not influence epidermal patterning in the genotypes analysed.

**Figure 3.**
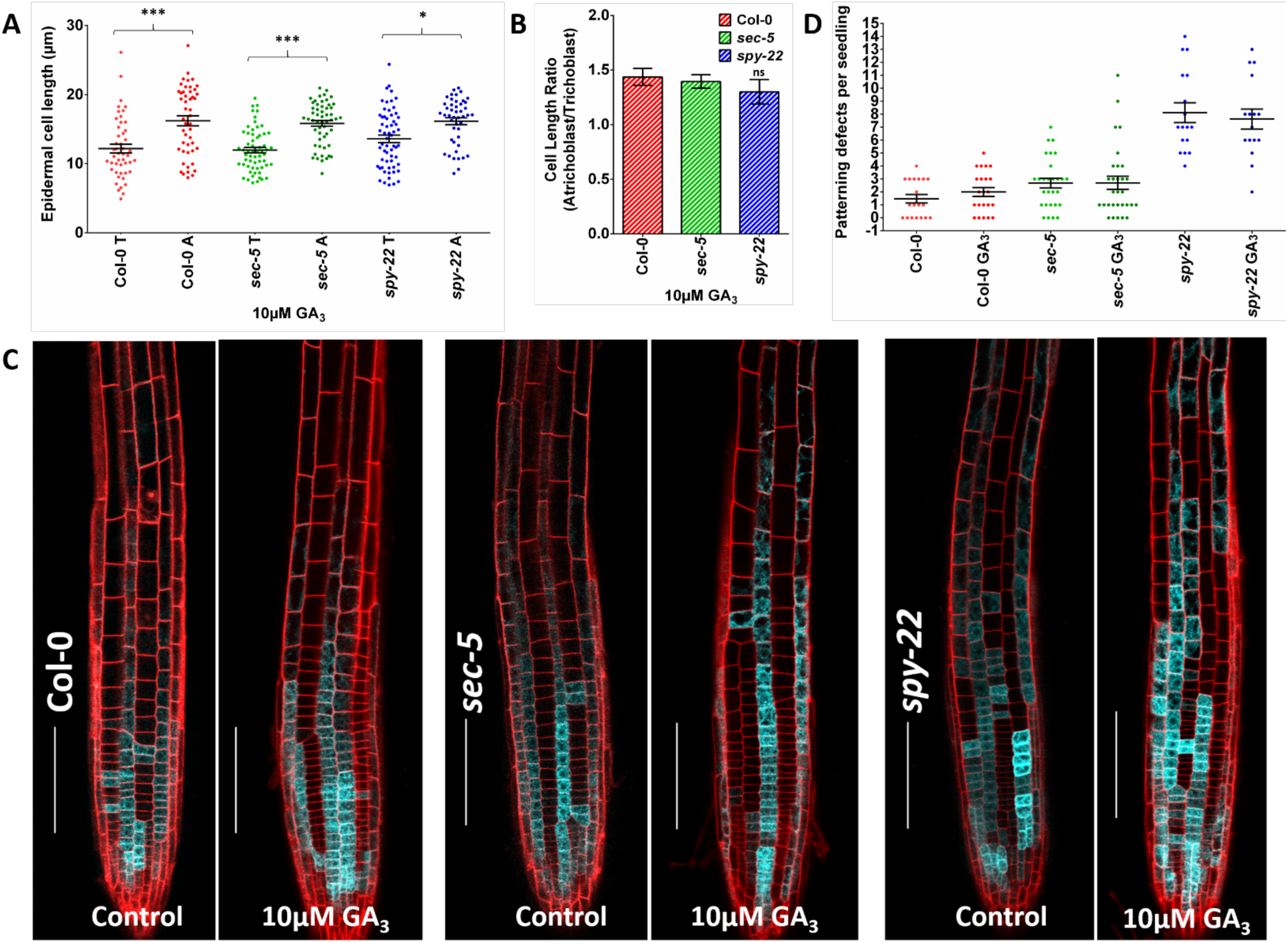
A- Epidermal cell length of 7-day old Col-0, *sec-5* and spy-22 seedlings grown on ½ MS supplemented with 10 µM GA_3_, n = 48-60. B- Presence of 10 µM GA_3_ does not influence the epidermal patterning, the ratio of the epidermal cell lengths of atrichoblasts/trichoblasts is lower in *spy-22* compared to *sec-5* and Col-0. C- GL2::4xYFP expression pattern remains largely unchanged in presence of 10 µM GA_3_, scale bar – 100µm. D- Patterning defects per seedling defined as the number of times an atrichoblast appears in trichoblast cell file and vice versa. The average number of patterning events per seedling remained unaffected in the presence of 10µM GA_3_ in all the lines compared to untreated controls. For statistical analysis, One-way ANOVA with Tukey’s multiple comparison was done (*** P ≤ 0.001, * P ≤ 0.05), data from three biological repeats is shown.

Gibberellin signalling in Arabidopsis is regulated via its ability to mediate the degradation of DELLA proteins, a family of growth inhibitors. The degradation of DELLAs de-represses the DELLA interacting proteins which in turn positively regulate growth (Bao *et al.* 2020; Davière and Achard, 2016). Most of the available literature on DELLAs is based on work in the L*er*-background. In order to mimic an environment with reduced GA signalling also in our mutant lines in Col-0 background, we deleted 17 amino acids of the DELLA domain of RGA as described by (Dill *et al.* 2001), preventing its recognition by the GA receptor GID. This resulting *RGA::*Δ*RGA* construct was transformed into Col-0, rendering the transformants insensitive to GA and thus constitutively repressing the DELLA interacting proteins. The resulting plant lines displayed similar phenotypes like described before in the L*er* background, including smaller leaf and rosette size, darker leaves, and reduced inflorescence axis length (Figure S3). We then crossed this line into *sec-5* and *spy-22*, in order to test whether reduced GA signalling impacts on ectopic root hair formation. Examination of *RGA::*Δ*RGA* Col-0 roots demonstrated that root hair patterning is similar to that of Col-0, showing no discernible ectopic root hair formation. *RGA::*Δ*RGA sec-5* and *RGA::*Δ*RGA spy-22* root meristems were indistinguishable from their *sec-5* and *spy-22* parents, respectively, with *RGA::*Δ*RGA spy-22* still displaying ectopic root hairs (Figure 4 A).

**Figure 4.**
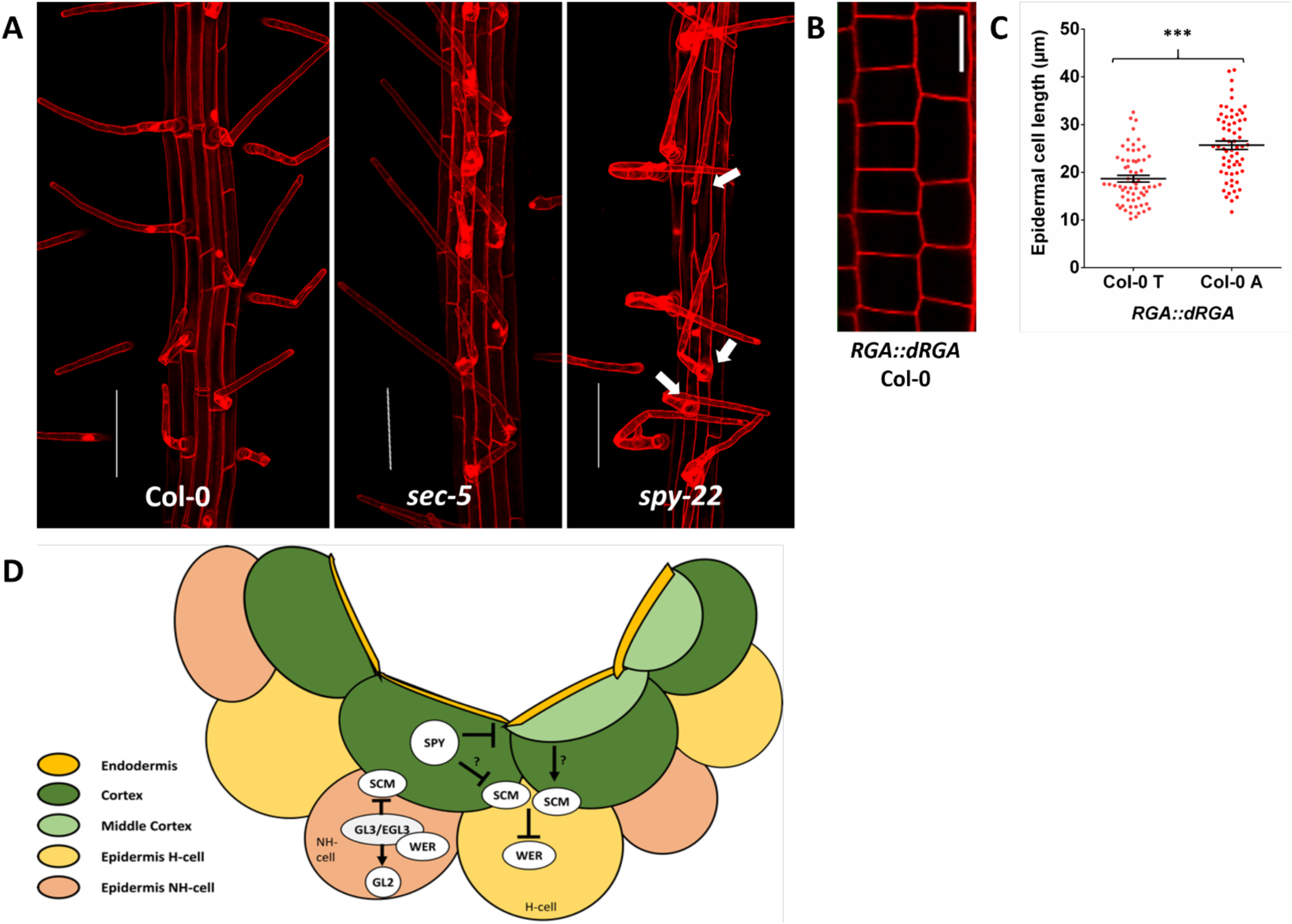
A- 7-day old *RGA::dRGA* Col-0, *RGA::dRGA sec-5* and *RGA::dRGA spy-22* seedlings grown on ½ MS agar mounted in PI. *RGA::dRGA* Col-0 and *RGA::dRGA sec-5* did not show ectopic root hairs, while in *RGA::dRGA spy-22* ectopic root hair formation was comparable to *spy-22* (see Figure 2 A), scale bar – 100µm. B- Epidermal layer of 7 day old *RGA::dRGA* Col-0 seedling in the late meristematic region. The length of 4 consecutive cells in neighbouring tricho- and atrichoblast files were measured, scale bar – 20µm. C- In *RGA::dRGA* Col-0, atrichoblasts cells are significantly larger than trichoblast cells, similar to Figure 1 D and 3 A. For statistical analysis a Students T-test.was done (*** P ≤ 0.001). D- Model describing the role of SPY in root hair patterning. H-cell: root hair cell/trichoblast. NH-cell: non-root hair cell/atrichoblast.

Above experiments suggest that epidermal cell patterning defects and ectopic root hair formation in *spy-22* are independent of GA signalling. Further, we measured the cell size of tricho- and atrichoblasts in the transition zone of *RGA::*Δ*RGA* Col-0 root meristems (Figure 4 B) and observed that the ratio between the two cell types was unaltered when compared to values obtained in the Col-0 parent background (Figure 4 C, compare with Figure 1 C, D). These findings suggest that epidermal cell patterning and differentiation in wild type roots is independent of GA signalling.

## Discussion

Root hairs are essential for the uptake of water and nutrients, as they can sense nutrients in the soil and react by increasing the root surface in a very flexible way. Root hair patterning is therefore regulated by internal as well as environmental factors, allowing for a high degree of plasticity in the developmental program. Thus, many different pathways feed into the regulation of cell fate determination in the epidermis, including a number of hormones such as auxin, ethylene and brassinosteroids (Balcerowicz *et al.* 2015; Borassi *et al.* 2020; Kuppusamy *et al.* 2009; Liu *et al.* 2018; Shibata and Sugimoto, 2019). Root hair patterning in Arabidopsis has been studied extensively and represents a very useful model system for analysis of plasticity in cell fate determination. In recent years, a number of tools have been made available to monitor the establishment of hair- and non-hair cell files in the root apical meristem, including a set of transcriptional reporters labelling specific cell types (Marquès-Bueno *et al.* 2016). Here, we present evidence that O-fucosylation is involved in establishing root hair cell patterning. Using a number of transcriptional reporters, genetics and phenotypical analysis, we show that root hair cell patterning is impaired in the O-fucosyltransferase mutant *spy-22*. Monitoring the expression of WER by using a transcriptional reporter suggests that the patterning defect in *spy-22* is established already early on during epidermal cell fate determination, potentially due to defects in cortex development or cell-to cell communication between cortex and epidermis, as these processes regulate cell type specific WER expression levels. The atypical receptor-like kinase SCRAMBLED (SCR) plays an important role in signalling from the cortex to the epidermis and further on to WER in this context (Gao *et al.* 2019; Kwak *et al.* 2005). Further experiments targeting the function, localization or turn-over of SCR might help determining how SPY participates in cell-to-cell communication at this stage, or alternatively in upstream signalling events in the cortex. Other potential targets of SPY include the transcription factor JACKDAW (JKD), that is expressed in the cortex and regulates epidermal cell fate in a non-cell autonomous way or other regulators of SCR, such as QKY (Hassan *et al.* 2010; Song *et al.* 2019).

Post-translational modification by attachment of O-fucose or O-GlcNAc is still not very well understood in plants. The best studied target is the gibberellin signalling repressor RGA, where O-GlcNAc and O-fucose have opposite effects on its activity, probably by inducing conformational changes (Zentella *et al.* 2016; Zentella *et al.* 2017). Accordingly, *spy*-mutants show many phenotypes that can be associated with gibberellin signalling, such as paclobutrazol resistance, early flowering, or elongated growth (Olszewski *et al.* 2010; Silverstone *et al.* 2007). In our study, we did not find an indication that consequences of altered O-fucosylation on root hair-patterning would require gibberellin signalling, as exogenous application of GA did not affect patterning (Figure 3). Consistently, we did not observe root hair patterning defects in *RGA::*Δ*RGA* lines (Figure 4). The observed increase in cell numbers of *spy-22* meristems (Figure S1) is probably independent of the patterning defect, but further studies are necessary to address if this increased cell division is dependent on GA-signalling.

Overall, we suggest a model, where SPY regulates root hair cell fate determination by affecting the spatial order of WER-expression, which then signals down to patchy expression of GL2 and EXP7, leading to ectopic root hair formation (Figure 2). Thus, O-glycosylation potentially regulates the function of upstream regulators such as SCM or the cell-to-cell communication from cortex to the epidermis (Figure 4 D), but further studies are necessary to reveal the direct targets of SPY in this context.

## Methods

### Plant material and growth conditions

All mutant lines used in this study were obtained from the Nottingham Arabidopsis Stock Centre NASC. Col-0 ecotype of *Arabidopsis thaliana* is referred to as wild-type control. T-DNA insertion lines of *spy-22* (SALK_090582) and *sec-5* (SALK_034290) and previously published reporter lines WER::4xYFP (N2106117), GL2::4xYFP (N2106121) and EXP7::4xYFP (N2106118) (Marquès-Bueno *et al.* 2016) in Col-0 background were used. After surface sterilisation with 70% ethanol, the seeds were plated onto half Murashige and Skoog medium (2.15 g/L MS Salts, 0.25 g/L MES, pH 5.7, 1% agar). After stratification in the dark at 4°C for 2 days, they were vertically grown in long day conditions (16 hours light / 8 hours dark) at 22°C.

### Microscopy

For imaging, a Leica TCS SP5 confocal microscope with an HCX PL APO CS 20.0×0.70 IMM UV objective was used. Seedlings were mounted in Propidium iodide (PI) (0.02 mg/mL) for staining the cell wall prior imaging. DPSS561 Laser was used to excite PI at 561nm (emission 584-735nm with standard PMT), and an Argon Laser at 30 % intensity was used to excite YFP at 514nm (emission 524-552 with HyD detector). Z Stacks were taken for visualizing root hairs and Maximum Projections were made using the Leica LAS AF lite software.

### Phenotyping and Image quantification

Measurements and quantifications were performed using the LAS X Leica Software. For studying the RAM length, seedlings were mounted in PI (0.02 mg/mL). We measured the distance from quiescent centre till the uppermost first cortical cell which was twice as long as wide as described by (Feraru *et al.* 2019). For epidermal cell patterning, lengths of 4 consecutive cells from neighbouring (tricho/atrichoblast) files in the late meristem were measured (Lofke *et al.* 2015). For analysing the patterning frequency in GL2::4xYFP, we checked for its expression in cell division and transition zones. We defined the occurrence of trichoblast cells in an atrichoblast cell file and vice versa as a patterning defect and counted the number of such patterning events in each seedling.

### Data Analysis

We used GraphPad Prism 5 and 6 for generating graphs. Error bars in graphs indicate standard error. One-way ANOVA and Tukey’s Multiple comparison test were performed for statistical analysis of the data. Sample sizes (n) for all experiments are given in the respective figure legends.

### Plasmid construction and generation of transgenic lines

To generate a GA insensitive, stabilized version of RGA in the Col-0 background, *RGA::dRGA* was amplified from genomic DNA of Col-0 using Q5 high fidelity DNA polymerase (NEB). Two overlapping fragments lacking 17 aminoacids covering the DELLA domain like described in (Feng *et al.* 2008) were generated using the following primer pairs: #270 (5’-tacaaaaaagcaggctccactagtactaattattcgtctgtc-3’) and #272 (5’-gttcgagtttcaaagcaacctcgtccatgttacctccaccgtc-3’), #273 (5’-gacggtggaggtaacatggacgaggttgctttgaaactcgaac-3’) and #271 (5’-gctgggtctagatatctcgagtacgccgccgtcgagag-3’); The resulting overlapping fragments were then cloned into a Gateway™ pENTR4™ vector backbone linearized with NcoI/XhoI via Gibson Assembly (NEB). The assembled plasmid was transformed into electrocompetent DH10b *E.coli* cells, positive clones were selected on LB medium using kanamycin (50µg/mL) and confirmed by sequencing. Confirmed entry clones were digested with AsiI to destroy the kanamycin resistance of the pENTR4-backbone, and recombined with pEarleyGate303 (Earley *et al.* 2006) using Gateway LR Clonase ll enzyme mix to generate a plant expression vector. Positive colonies were selected for kanamycin (50µg/mL) resistance, confirmed plasmids were electro-transformed into *Agrobacterium tumefaciens* GV3101 and used for transforming *Arabidopsis thaliana* ecotype Col-0 by floral dipping (Clough and Bent, 1998). Stable transformants with a strong GA-deficient phenotype were selected before crossing with *spy-22* and *sec-5*.

## Acknowledgements

We are grateful to Monika Debreczeny, Barbara Korbei, Jürgen Kleine-Vehn, and members of his group for numerous discussions and support with setting up microscopy techniques, and Mathias Ried for technical support. We thank Christian Luschnig and Melina Velasquez for critically reading the manuscript. Funding was provided by the Austrian Academy of Sciences ÖAW (DOC-fellowship to KVM, APART fellowship to DL) and the Austrian Science Fund FWF (Project number P20051).

## Author contributions

KVM and DL planned experiments, IZ provided substantial technical support, KVM wrote the manuscript with support by DL.

---

**Supplement Figure 1.**
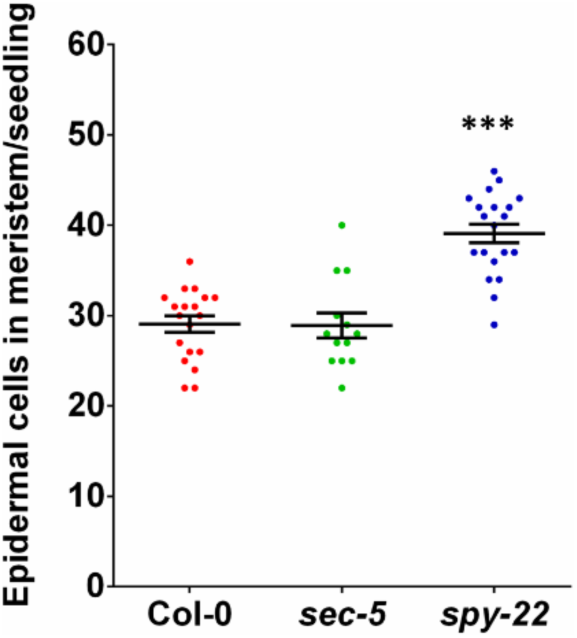
Number of epidermal cells in the meristem of 7 DAG O-glycosylation mutants. Meristem of *spy-22* mutants have a higher number of epidermal cells (39.10 +/- 4.599) compared to Col-0 (29.05 +/- 3.965) and *sec-5* (28.92 +/- 5.008). For statistical analysis, One-way ANOVA with Tukey’s multiple comparison was done (*** P ≤ 0.001), data from three independent biological repeats is shown.

**Supplement Figure 2.**
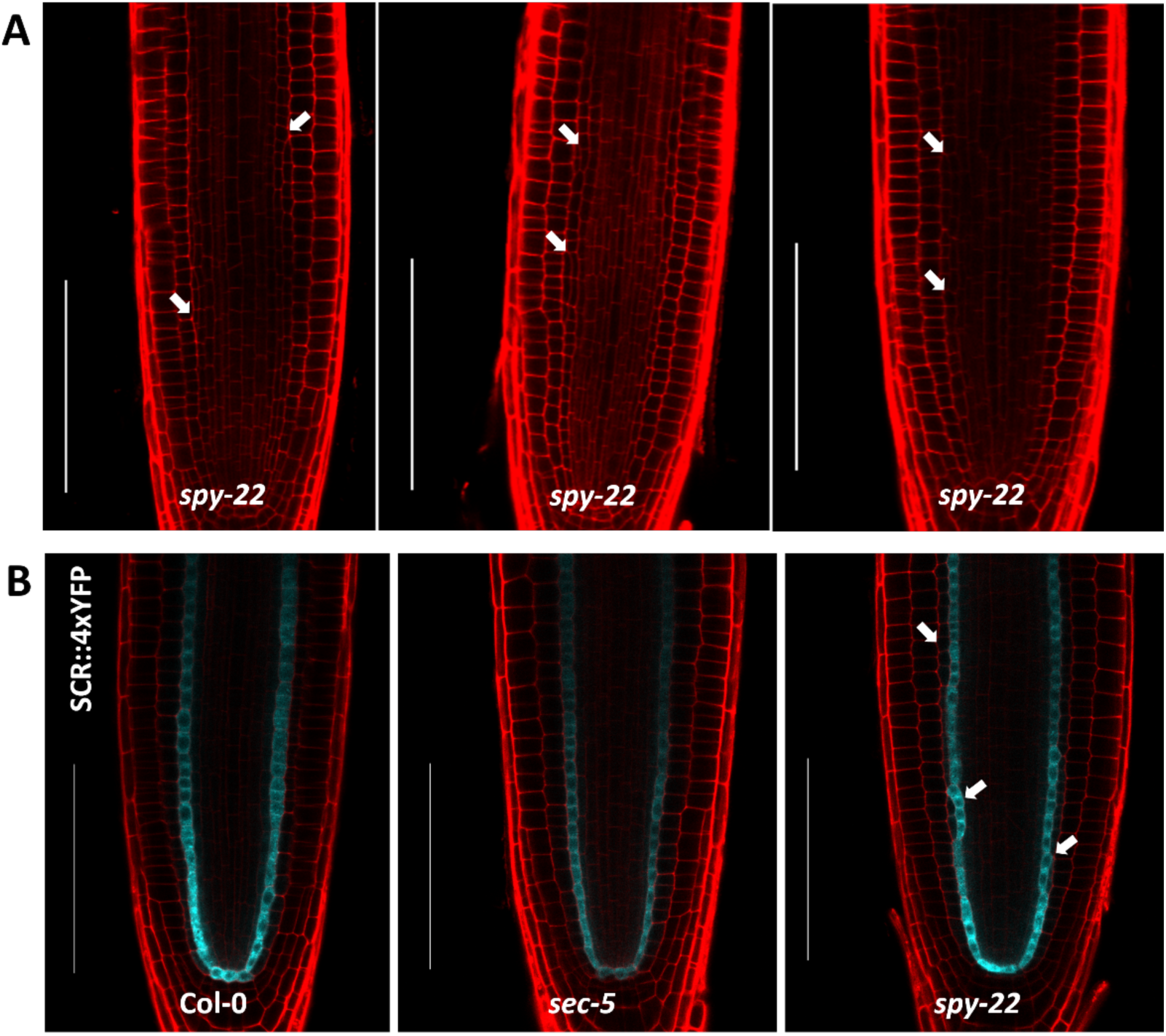
A – 7-day old *spy-22* seedlings grown on ½ MS agar mounted in PI, arrows indicate middle cortex formation. This extra layer of cortex is formed between cortex and endodermis and has been previously described by Cui *et al.* 2014. scale bar – 100µm. B - SCR::4xYFP expression in Col-0, *sec-5* and *spy-22* is restricted to the endodermis. The middle cortex proliferation in the *spy-22* background is unique and independent of SCR expression in the endodermis.

**Supplement Figure 3.**
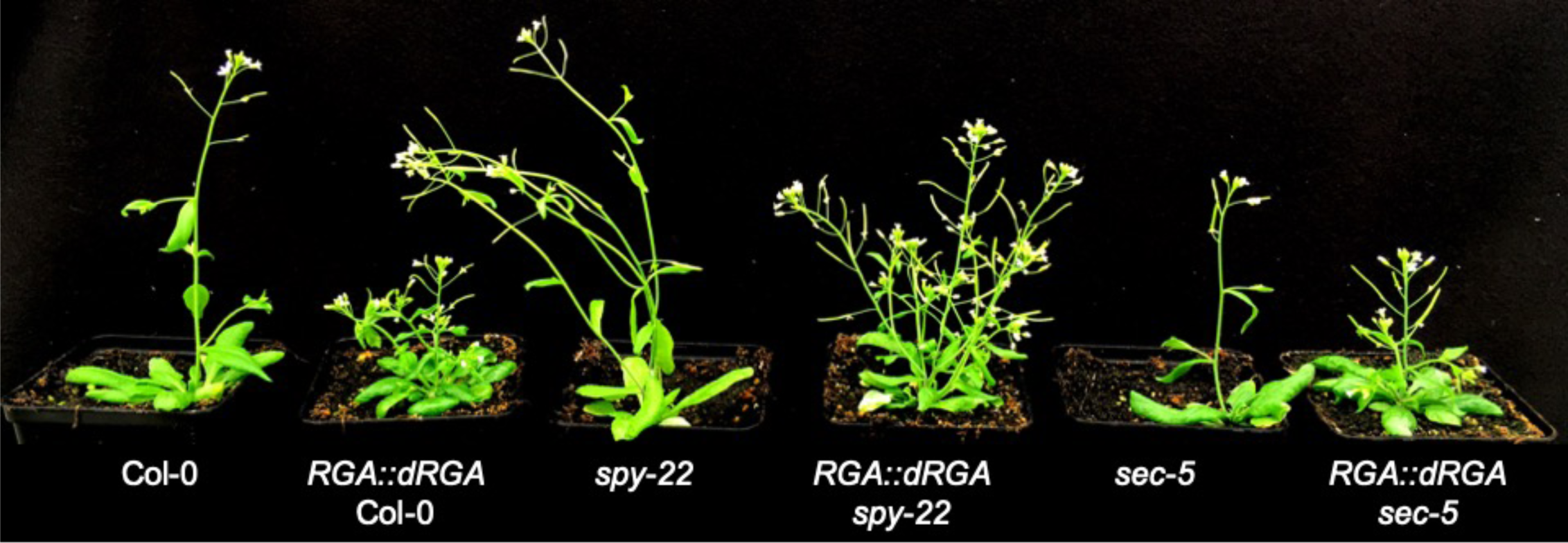
Col-0, *spy-22, sec-5* and their crosses with *RGA::dRGA* Col-0, a line expressing a stabilized version of the GA-signaling repressing DELLA protein RGA. All *RGA::dRGA* lines show phenotypes characteristic for low GA signaling, like smaller rosette size and shorter inflorescences.

